# Inferring interaction partners from protein sequences

**DOI:** 10.1101/050732

**Authors:** Anne-Florence Bitbol, Robert S. Dwyer, Lucy J. Colwell, Ned S. Wingreen

## Abstract

Specific protein-protein interactions are crucial in the cell, both to ensure the formation and stability of multi-protein complexes, and to enable signal transduction in various pathways. Functional interactions between proteins result in coevolution between the interaction partners. Hence, the sequences of interacting partners are correlated. Here we exploit these correlations to accurately identify which proteins are specific interaction partners from sequence data alone. Our general approach, which employs a pairwise maximum entropy model to infer direct couplings between residues, has been successfully used to predict the three-dimensional structures of proteins from sequences. Building on this approach, we introduce an iterative algorithm to predict specific interaction partners from among the members of two protein families. We assess the algorithm's performance on histidine kinases and response regulators from bacterial two-component signaling systems. The algorithm proves successful without any *a priori* knowledge of interaction partners, yielding a striking 0.93 true positive fraction on our complete dataset, and we uncover the origin of this surprising success. Finally, we discuss how our method could be used to predict novel protein-protein interactions.

## INTRODUCTION

Many key cellular processes are carried out by interacting proteins. For instance, transient protein-protein interactions determine signaling pathways, and their specificity ensures proper signal transduction. Hence, mapping specific protein-protein interactions is central to a systems-level understanding of cells, and has broad applications to areas such as drug targeting. High-throughput experimental methods have recently elucidated a substantial fraction of protein-protein interactions in a few model organisms [1], but experimental approaches remain challenging. Meanwhile, major progress in sequencing has led to an explosion of available sequence data. Can we exploit this abundant new sequence data to identify specific protein-protein interaction partners?

Specific interactions between proteins imply evolutionary constraints on the interacting partners. For instance, mutation of a contact residue in one partner generically impairs binding, but may be compensated by a complementary mutation in the other partner. This co-evolution of interaction partners results in a correlation of their amino-acid sequences. Similar correlations exist within single proteins, between amino acids that are in contact in the folded protein. However, the simple fact of a correlation between residues in a multiple sequence alignment is only weakly predictive of a three-dimensional contact [2–4], as correlation can also stem from other effects such as phylogeny and indirect interactions. Fortunately, global statistical models provide a means to disentangle direct and indirect interactions [5–7]. In particular, the maximum entropy principle [8] specifies the least-structured global statistical model consistent with the one- and two-point statistics of an alignment [5]. This approach has recently been used with success to determine three-dimensional protein structures from sequences [9–11], to predict mutational effects [12–14], and to find residue contacts between known interaction partners [7, 15–19]. Pairwise maximum entropy models have also been used productively in various other fields (see e.g. Refs. [20–25]).

Here we present a pairwise maximum entropy approach that uses sequence data to predict specific interaction partners among the paralogous genes belonging to two families of interacting proteins. We use histidine kinases (HKs) and response regulators (RRs) from prokaryotic two-component signaling systems (Fig. 1A) as a benchmark. Two-component systems constitute a major class of transduction pathways that enable bacteria (and archaea) to sense and respond to various signals from their environment. Typically, a transmembrane HK senses a signal, autophosphorylates, and transfers its phosphate group to its cognate RR protein, which induces a cellular response [27]. Most HKs are encoded in operons together with their cognate RR, so specific interaction partners are known, which enables us to assess performance. There are often dozens of paralogs of HKs and RRs within each genome, making the task of predicting cognate interaction partners from sequences alone highly nontrivial.

**FIG. 1.**
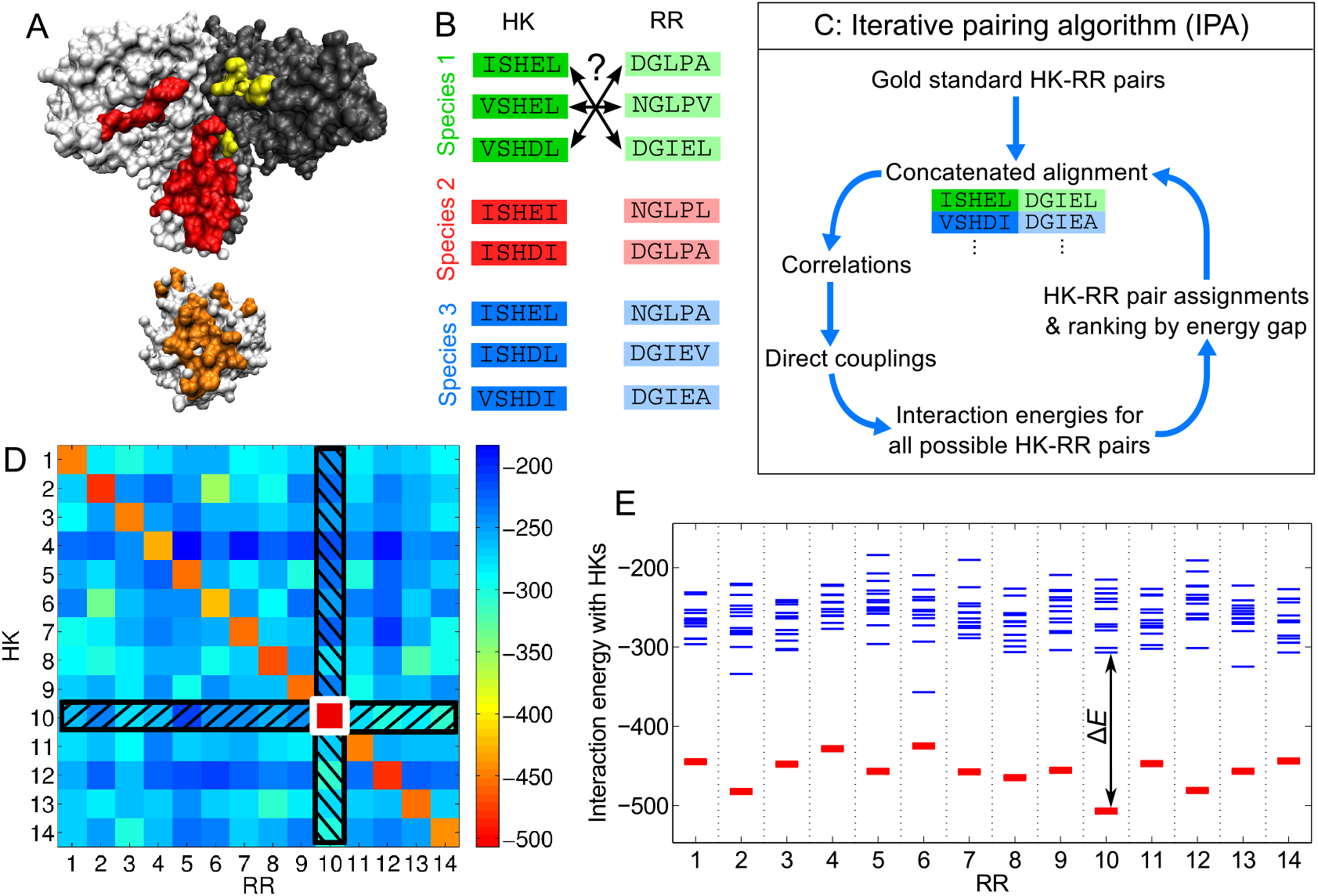
Iterative pairing algorithm (IPA). (A) Surface representations of a histidine kinase dimer (HK, top) and a response regulator (RR, bottom), from a co-crystal structure [26]; the HK-RR contacts in each molecule are highlighted in color. (B) To correctly pair HKs and RRs in each species from their sequences alone, we start from multiple sequence alignments of HKs and RRs, including 64 amino acids from the HK and 112 from the RR. (C) Schematic of the main steps of the IPA. (D, E) Example of HK-RR pair assignment and ranking by energy gap for one species. (D) Color map of the matrix of HK-RR interaction energies in *E*. *coli* K-12 MG1655 from the final iteration of the IPA performed on our standard dataset, with a gold standard set of *N*_start_ = 100 HK-RR pairs, and an increment step of *N*_increment_ = 200 pairs. As in every IPA iteration and every species, the pair with the lowest interaction energy is selected first (here, HK 10 - RR 10, boxed in white), and this HK and RR are removed from further consideration (black hatches). Then, the next pair with the lowest energy is chosen, and the process is repeated until all HKs and RRs are paired. (E) Energy spectrum from (D), showing the interaction energies with all the HKs for each RR, with the correct HK-RR pairs shown in red. The energy gap Δ*E* is shown for the RR with the highest gap (RR 10). A confidence score based on the energy gap is used to rank all assigned HK-RR pairs, in order to build the concatenated alignment for the subsequent IPA iteration. See Materials and Methods for details.

To address this challenge, we developed an iterative pairing algorithm (IPA, Fig. 1). At each iteration, the highest-scoring predicted HK-RR pairs are incorporated into the concatenated sequence alignment from which we build the pairwise maximum entropy model. We show that this approach yields a major increase of predictive accuracy through progressive training of the model. First, we consider the case where the IPA starts with a set of known HK-RR partners. We obtain good performance even starting from small numbers of known pairs. Then, we show that the IPA can make accurate predictions *starting without any known pairings*, as would be needed to predict novel protein-protein interactions. We trace the origin of this success to the preferential recruitment of new correct pairs by the true pairs already in the concatenated alignment. Finally, we discuss how multiple random initializations can be leveraged to improve the predictive accuracy of the IPA, and ultimately to identify new interactions between protein families.

## RESULTS

### Iterative pairing algorithm

We have developed an iterative method to predict interaction partners among the paralogs of two protein families in each species, just from their sequences (Fig. 1A,B). In each iteration (Fig. 1C; Materials and Methods), we compute correlations between residues from a concatenated alignment (CA) of paired sequences. The initial CA can either be built from a “gold standard” set of correct protein pairs that are assumed to be known, or be made from random pairs, assuming no prior knowledge of interacting pairs. We then infer the direct couplings for all residue pairs using a pairwise maximum entropy model of the CA and a mean-field approximation [9, 28]. We calculate the interaction energy for every possible protein pair within each species, by summing the inter-protein couplings. Note that these “energies” capture evolutionary correlations but cannot be interpreted directly as physical energies, though they correlate to energies for lattice proteins [29]. Using these interaction energies, we predict pairs (under the well-supported assumption of one-to-one specific HK-RR interactions [27], Fig. 1D). For each predicted pair, we compute a confidence score based on the energy gap between the pair with the lowest interaction energy and the next alternative pair (Fig. 1E). The CA is then updated by including the protein pairs that have the highest confidence scores, and the next iteration can begin.

Unless otherwise specified in what follows, our results were obtained on a standard dataset comprising 5064 HK-RR pairs for which the correct pairings are known from gene adjacency. Each species has on average 〈*m_p_*〉 = 11.0 pairs, and at least two pairs, to avoid trivial cases (see Materials and Methods).

### Starting from known pairings

We begin by predicting interaction partners starting from a “gold standard” set of known pairs. To implement this, we pick a random set of *N*_start_ known HK-RR pairs from our standard dataset, and the first IPA iteration uses this concatenated alignment (CA) to train the model. We blind the pairings of the remaining sequences and use them as our “test” set, on which we predict pairings. At each subsequent iteration *n* > 1, the CA contains the (*n* – 1)*N*_increment_ highest-scoring pairs from the previous iteration in addition to the gold standard pairs (see Materials and Methods), and the model is retrained on this larger dataset.

At the first iteration, the fraction of accurately predicted HK-RR pairs (TP fraction) depends strongly on *N*_start_, and is close to the random expectation (0.09) for very small gold standard sets, ranging from 0.13 at *N*_start_ = 1 to 0.93 for *N*_start_ = 2000 (Fig. 2, inset, blue curve). With subsequent iterations the TP fraction steadily increases (Fig. 2, main panel). Strikingly, the final TP fraction depends only weakly on *N*_start_, remaining high even for small gold standard sets. For *N*_start_ = 1, the IPA achieves a final TP fraction of 0.84, a huge increase from the initial value of 0.13 (Fig. 2, inset, red curve). This result demonstrates the power of the iterative approach, in which adding high-scoring pairs at each iteration progressively increases the predictive accuracy of the model.

**FIG. 2.**
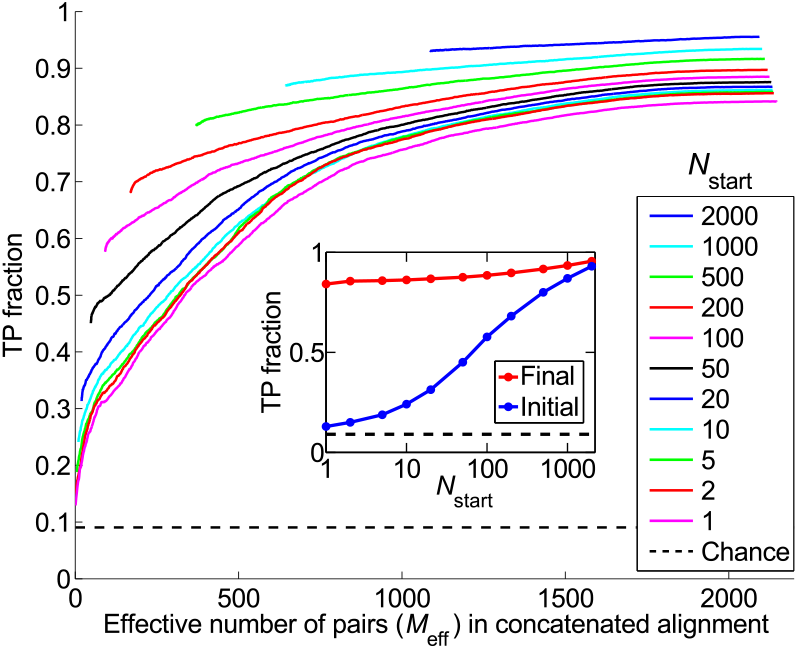
Fraction of predicted pairs that are true positives (TP fraction), for different gold standard set sizes *N*_start_. Main panel: Progression of the TP fraction during iterations of the IPA. The TP fraction is plotted versus the effective number of HK-RR pairs (*M*_eff_, see Supporting Information, Eq. S1) in the concatenated alignment, which includes *N*_increment_ = 6 additional pairs at each iteration. The IPA is performed on the standard dataset, and all results are averaged over 50 replicates that differ by the random choice of pairs in the gold standard set. Dashed line: Average TP fraction obtained for random HK-RR pairings. Inset: Initial and final TP fractions (at first and last iteration) versus *N*_start_, from the same data.

### Starting without known pairings

Given the success of the IPA with very small gold standard sets, we next ask whether predictions can be made without assuming any knowledge of interacting pairs. To test this, we randomly pair each HK with an RR from the same species, and use these 5064 random pairs to train the initial model. At each subsequent iteration *n* > 1, the CA is built just from the (n – 1)*N*_increment_ highest-scoring pairs from the previous iteration (see Supporting Information).

Fig. 3 shows the progression of the TP fraction for different values of *N*_increment_. In all cases, the TP fraction increases steadily, with the increase being larger for smaller *N*_increment_. Thus, our iterative method works best when the increment step is small (Fig. 3, inset). The low–*N*_increment_ limit of the final TP fraction is 0.84, identical to that obtained with a single correct training pair (Fig. 2). We emphasize that this striking TP fraction of 0.84 is attained without any a priori knowledge of HK-RR interactions: the IPA allows us to bootstrap our way toward high predictivity. The low–*N*_increment_ limit is almost reached for *N*_increment_ = 6; thus we generically use *N*_increment_ = 6 to reduce computational time while retaining near-optimal performance. Note that the final TP fraction is robust with respect to different initializations: for *N*_increment_ = 6, its standard deviation, calculated on 500 replicates, is 0.04.

**FIG. 3.**
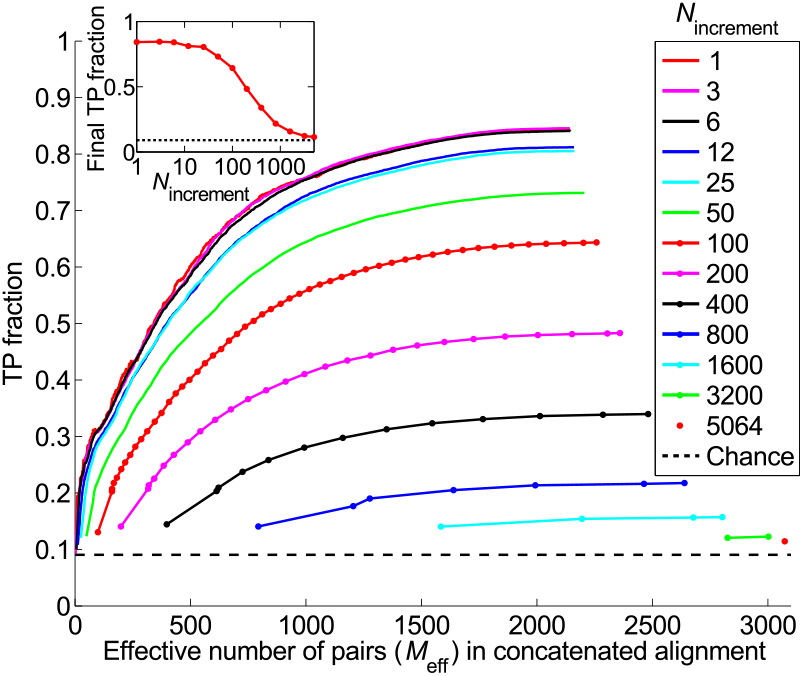
Starting from random pairings, i.e. without known pairings. Main panel: TP fraction plotted during iterations of the IPA versus the effective number of HK-RR pairs (*M*_eff_) in the concatenated alignment, which includes *N*_increment_ additional pairs at each iteration. Different curves correspond to different values of *N*_increment_. The IPA is performed on the standard dataset, and all results are averaged over 50 replicates that differ in their initial random pairings. Note that the first point of each curve corresponds to the second iteration. Dashed line: Average TP fraction obtained for random HK-RR pairings. Inset: Final TP fraction versus *N*_increment_, from the same data.

### Training process

The ability to accurately predict interaction partners without training data is surprising. To understand it, we examine the evolution of the model over iterations of the IPA. In a well-trained model, the residue pairs with the largest normalized couplings have been shown to correspond to contacts in the protein complex [7, 16, 30]. Up to iteration ~100 – 150 (with *N*_increment_ = 6), models starting from random pairings do no better than chance at identifying contacts. Above this number of iterations, the models improve rapidly and soon predict contacts nearly as well as models constructed from the same number of correct HK-RR pairs (Figs. 4 and S1A).

**FIG. 4.**
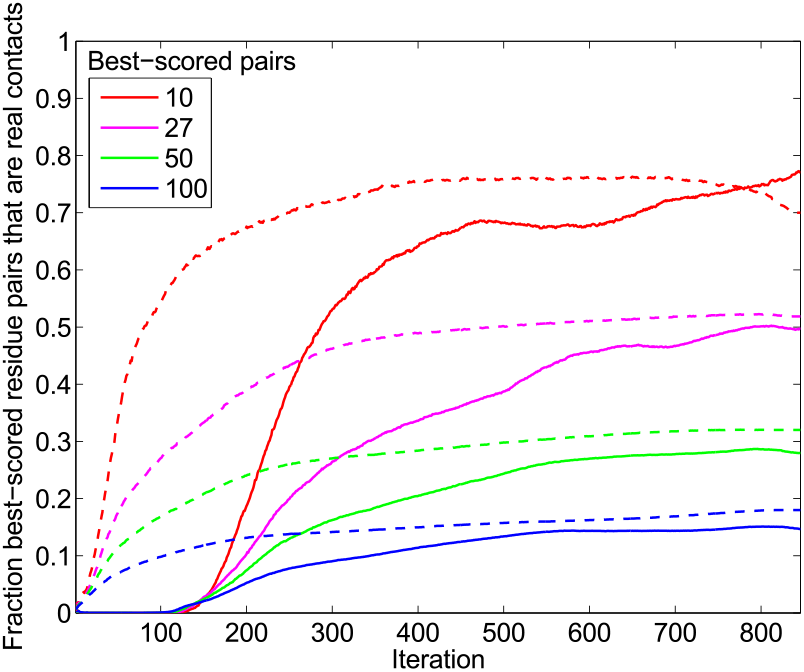
Training of the couplings during the IPA. Residue pairs comprised of an HK site and an RR site are scored by the Frobenius norm (i.e. the square root of the summed squares) of the couplings involving all possible residue types at these two sites. The best-scored residue pairs are compared to the 27 HK-RR contacts found experimentally in Ref. [26]. Solid curves: Fraction of residue pairs that are real contacts (among the *k* best-scored pairs for four different values of *k*) versus the iteration number in the IPA. Dashed curves: Ideal case, where at each iteration *N*_increment_ randomly-selected correct HK-RR pairs are added to the CA. The overall fraction of residue pairs that are real HK-RR contacts, yielding the chance expectation, is 3.8 × 10^-3^. The IPA was performed on the standard dataset with *N*_increment_ = 6, and all data is averaged over 500 replicates that differ in their initial random pairings.

At early stages, the model, constructed from a few rather randomly selected HK-RR pairs, is a poor predictor of real contacts, and of correct HK-RR pairs. However, couplings associated to residue pairs that occur in the CA are increased, raising the scores of HK-RR pairs with high sequence similarity to those already in the CA, and making them more likely to be added to the CA. With this in mind, we examine the degree of sequence similarity between the HK-RR pairs in the CA in consecutive iterations. As a concrete measure, we consider two HK-RR pairs to be “neighbors” if the sequence identity between the two HKs and between the two RRs are both > 70%. We find that sequence similarity is crucial in the recruitment of new HK-RR pairs to the CA at early iterations (Fig. S1B).

Understanding the initial increase of the TP fraction requires a further observation. In our standard dataset, among all possible, within-species HK-RR pairs, the average number of neighbor pairs per correct HK-RR pair is 9.66, of which 99% are correct. In contrast, the average number of neighbor pairs per incorrect HK-RR pair is 5.25, of which less than 1% are correct. In other words, correct pairs are more similar to each other than they are to incorrect pairs, or than incorrect pairs are to each other. We call this the *Anna Karenina effect*, in reference to the first sentence of Tolstoy’s novel [31]: All happy families are alike; each unhappy family is unhappy in its own way. Biologically, this makes sense: each HK-RR pair is an evolutionary unit, so a correct pair is likely to have orthologs of both the HK and the RR in multiple other species, whereas an incorrect pair is less likely to have orthologs of both the HK and the (non-cognate) RR in other species. Hence, in early iterations, the number of neighbors recruited per correct pair is significantly higher than that per wrong pair (Fig. S1B), increasing the TP fraction in the CA. To summarize, sequence similarity is crucial at early stages in the bootstrapping process, and the Anna Karenina effect helps to increase the TP fraction in the CA, ultimately promoting training of the model. (Thus the IPA might be further enhanced by an initial phase where HK-RR pairs are scored based on similarity [32].)

### Dependence on features of the dataset

To apply the IPA approach more widely, it is important to understand what characteristics of the dataset enable its success. The number of pairs per species is likely important, since pairing is more difficult when there are more incorrect possibilities. Indeed, higher TP fractions are obtained in datasets with fewer average pairs per species (Fig. 5A), and the presence of species with a small number of pairs is crucial (Fig. S2). (In their absence, the TP fraction can be rescued by a sufficiently large set of known pairs (Fig. S3).) Perhaps surprisingly, for small *N*_increment_, the final TP fraction does not depend on how many pairs in the initial CA are correct (Fig. 5A, red curves, and Fig. S4). Hence, the importance of species with few pairs does not stem from a more favorable initialization. Rather, HK-RR pairs from species with few pairs tend to obtain higher confidence scores since they have fewer competitors, making them more likely to enter the CA at early stages (Fig. S5). This bias in favor of species with few pairs combines with the Anna Karenina effect to favor the recruitment of correct pairs to the CA early in the learning process.

**FIG. 5.**
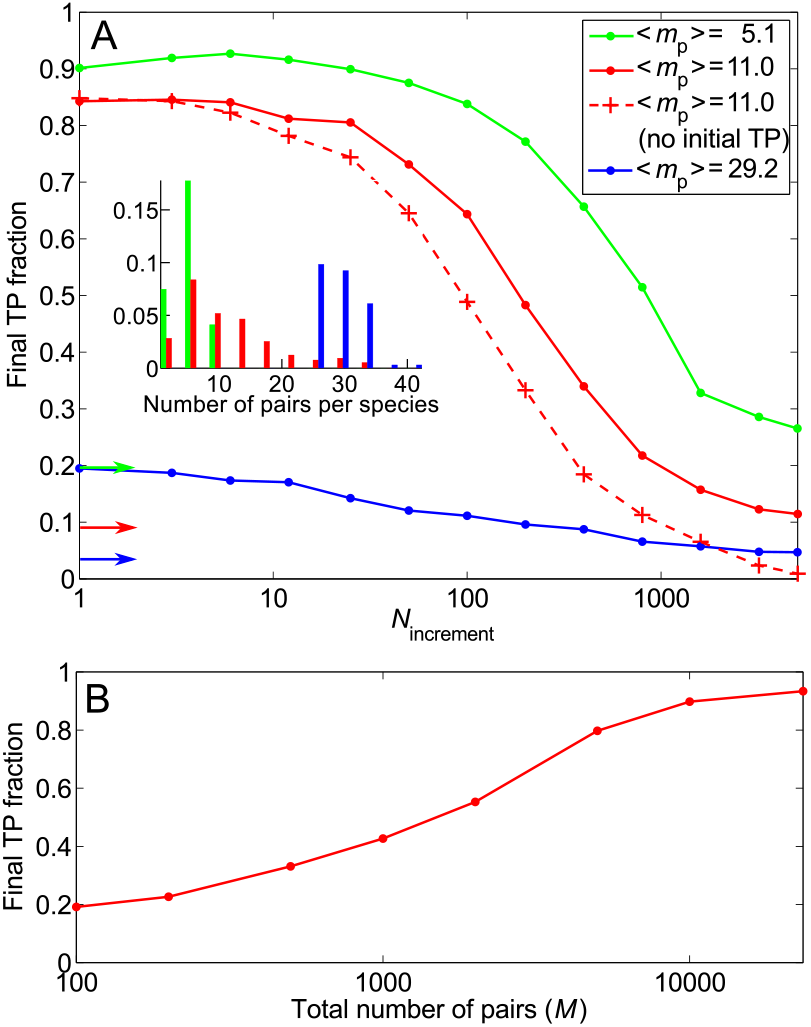
Impact of the number of HK-RR pairs per species, and of the total number of HK-RR pairs in the dataset. (A) Main panel: Final TP fraction versus *N*_increment_ for three datasets with the same total number of HK-RR pairs *M*, but with different distributions of the number of pairs per species yielding different means 〈*m_p_*〉. Solid curves: Starting from random pairings. Dashed curve: Starting from random pairings with no initial correct pairs. The IPA is performed on the standard dataset with *N*_increment_ = 6, all results are averaged over 50 replicates, and colored arrows indicate the average TP fractions obtained for random HK-RR pairings in each dataset. Inset: Distribution of the number of pairs per species in the different datasets (red: standard dataset; green and blue: datasets comprised of the species with lowest or highest numbers of pairs in the full dataset). (B) Final TP fraction versus the total number *M* of HK-RR pairs in the dataset, starting from random pairings. For each *M*, datasets are constructed by picking species randomly from the full dataset, preserving the average distribution of the number of HK-RR pairs per species. For each *M* except the largest, results are averaged over multiple different such alignments (from 50 up to 500 for small *M*). For the largest *M* (full dataset), averaging is done on 50 different initial random pairings. All results correspond to the small- *N*_increment_ limit.

Since sequence similarity is crucial at early iterations, it should strongly impact performance. Indeed, a lower final TP fraction (0.58 vs. 0.84) is obtained in a dataset where no two correct pairs are > 70% identical, but it can be rescued by a sufficiently large set of known pairs (Fig. S6).

A final important parameter is dataset size. The final TP fraction increases steeply above ~1000 sequences, and saturates above ~10, 000 (Fig. 5B). For the complete dataset (23,424 HK-RR pairs, see Materials and Methods), we obtain a striking final TP fraction of 0.93. Larger datasets imply closer neighbors, which is favorable to the success of bootstrapping. Moreover, in our case, ~500 correct pairs are needed for proper training (Fig. 2, inset, blue curve).

### Optimization

To improve the predictive ability of the IPA, we exploit multiple different random initializations of the CA. For each possible, within-species HK-RR pair, we calculate the fraction *f_r_* of replicates of the IPA in which this pair is predicted. High *f_r_* values are excellent predictors of correct pairs, significantly outperforming average TP fractions from individual replicates (Fig. 6, main panel). The quality of *f_r_* as a classifier is demonstrated by the area under the receiver operating characteristic: it is 0.991, very close to 1 (ideal). The strikingly high TP fraction of the pairs with highest *f_r_* values can be exploited by considering some of these pairs as a training set and running the IPA again. This “re-bootstrapping” process can be iterated, yielding further performance increases, particularly for small datasets (Fig. S7).

**FIG. 6.**
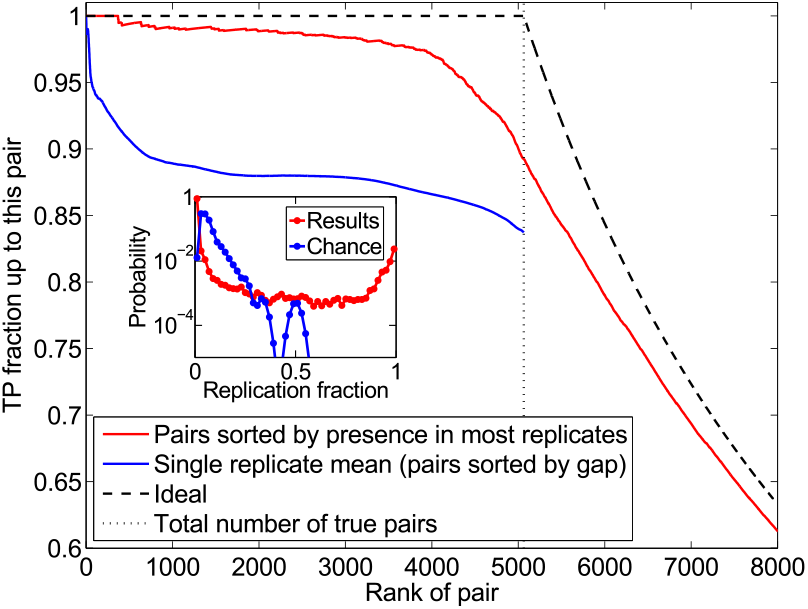
Improved accuracy from multiple initial random pairings. Red curve: All possible HK-RR pairs (within each species) are ranked by the fraction *f_r_* of replicates of the IPA in which they are predicted. The TP fraction up to each pair is plotted versus the rank of this pair. The standard dataset is used, with *N*_increment_ = 6. 500 replicates that differ in their initial random pairings are considered. Blue curve: For each separate replicate, pairs are ranked by their confidence score, in decreasing order. The TP fraction up to each pair is computed, and the mean of these curves is shown. Dashed curve: Ideal classification, where the *M* = 5064 first pairs (dotted line) are correct, while all the others are incorrect. Inset: Distribution of the fraction of replicates *f_r_* in which each possible HK-RR pair is predicted. Red curve: *f_r_* over 500 replicates of the IPA (same data as in main panel). Blue curve: Random HK-RR pairings within each species of the standard dataset.

### Toward predicting new protein-protein interactions

The distribution of replication fractions *f_r_* strongly favors values close to 0, mostly corresponding to wrong pairs, and close to 1, mostly corresponding to correct pairs (Fig. 6, inset). This is very different from the distribution expected for random within-species HK-RR pairings and from the distribution for a dataset lacking correct HK-RR pairs (Fig. S8). This characteristic bimodal-ity of *f_r_* for interacting proteins opens the way to using a bootstrapping method such as ours to test for specific interactions between members of two protein families.

## DISCUSSION

We have presented a method to infer specific interaction partners among the members of two protein families with multiple paralogs, using only sequence data. Our approach is based on pairwise maximum entropy models, which have proved successful at predicting residue contacts between known interaction partners [7, 15–19]. To our knowledge, the important problem of predicting interaction partners among paralogs from sequences has only been addressed by Burger and van Nimwegen [6], who used a Bayesian network method. Pairwise maximum entropy-based approaches were later shown to outperform this method for orphan HK-RR partner predictions, starting from a substantial training set of partners known from gene adjacency [15]. Our method is the first to address and solve the problem of predicting interaction partners without any initial known pairs, as even the seminal study [6] included a training set via species that contain only a single pair.

We benchmarked our iterative pairing algorithm (IPA) on prokaryotic HK-RR pairs. The top-scoring predicted HK-RR pairs are progressively incorporated into the concatenated alignment used to build the pairwise maximum entropy model. This allows for progressive training of the model, which yields major increases of predictive accuracy. Strikingly, the IPA is very successful even in the absence of any *a priori* knowledge of HK-RR interactions, yielding a 0.93 TP fraction on our complete HK-RR dataset. This bootstrapping process is aided most by species with few HK-RR pairs, and its success relies on initial recruitment of pairs by sequence similarity. In particular, correct pairs are more similar to one another than incorrect pairs, favoring recruitment of correct pairs - a process we called the “Anna Karenina effect”. Bootstrapping performance is best for datasets including many species and featuring strong sequence similarity, and when species with few pairs are included. The first two conditions are easily met for HK-RRs (a 0.84 TP fraction was obtained with 5064 paired sequences, out of 23,424 available, and our re-bootstrapping approach yields a 0.64 TP fraction even for a dataset of only 502 paired sequences from 40 species, Fig. S7B) and are realized for a large and growing number of other protein families. Regarding pairs per species, the HK-RR system actually constitutes a hard case [27]: most other protein families feature fewer paralogs per species.

Our approach could be combined with those of Refs. [7, 13, 15–19] to improve the prediction of residue contacts between protein partners. It solves the major problem [17–19] of finding the correct interaction partners among paralogs, which is a prerequisite for accurate contact prediction. In particular, better paralog-partner predictions will help extend accurate contact prediction to currently-inaccessible cases such as eukaryotic proteins, for which genome organization cannot be used to find interaction partners. Finally, our method paves the way toward the prediction of novel protein-protein interactions between protein families from sequence data alone.

## MATERIALS AND METHODS

Extended Materials and Methods are presented in the Supporting Information.

### Dataset

Our dataset was built from the online database P2CS [33, 34], which includes two-component system proteins from all fully-sequenced prokaryotic genomes. All data can thus be accessed online. We considered the protein domains through which HKs and RRs interact, more precisely the Pfam HisKA domain present in most HKs (64 amino acids) and the Pfam Response Reg domain present in all RRs (112 amino acids). We focused on proteins with known cognate partners, i.e. those containing only one such domain, and encoded in the genome in pairs containing an HK and an adjacent RR. Discarding species with only one pair, for which pairing is trivial, we obtained a complete dataset of 23,424 HK-RR pairs from 2102 species. A smaller “standard dataset” of 5064 pairs from 459 species was extracted by picking species randomly.

### Iterative pairing algorithm (IPA)

Here, we summarize each of the steps of an iteration of the IPA (Fig. 1C).

1. **Correlations**. Each iteration begins by the calculation of empirical correlations from the CA of paired HK-RR sequences. At each amino-acid site *i*, a given concatenated sequence can feature any amino acid (or a gap): these states are denoted by *α*. The empirical one- and two-site frequencies, *f_i_*(*α*) and *f_ij_*(*α*,*β*), of occurrence of amino-acid states are computed for the CA, using a re-weighting of similar sequences, and a pseudocount correction, to ameliorate issues due to biased sampling and to the finite size of the CA (Eqs. S1-S4) [7, 9, 15, 28]. The correlations are then computed as

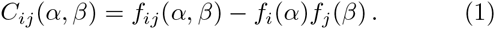
2. **Direct couplings**. Next, we construct a pairwise maximum entropy model of the CA (Eq. S6). It involves one-body fields *h_i_* at each site *i* and (direct) couplings *e_ij_* between all sites *i* and *j*, which are determined by imposing consistency of the global model with the empirical one- and two-point frequencies of the CA (Eqs. S7-S8). We use the mean-field approximation [9, 28] to solve this inference problem, i.e. the couplings are obtained by inverting the matrix of correlations:

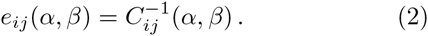 We then transform to the zero-sum gauge [7, 30].
3. **Interaction energies for all possible HK-RR pairs**. The interaction energy *E* of each possible HK-RR pair within each species of the dataset is calculated as a sum of direct couplings:

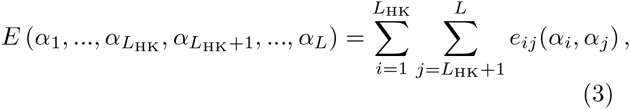 where *L*_HK_ denotes the length of the HK sequence and *L* that of the concatenated HK-RR sequence.
4. **HK-RR pair assignments and ranking by gap**. In each separate species, the pair with the lowest interaction energy is selected first, and the HK and RR from this pair are removed from further consideration, assuming one-to-one HK-RR matches (Fig. 1D). Then, the next pair with the lowest energy is chosen, and the process is repeated until all HKs and RRs are paired. Each assigned HK-RR pair is scored at assignment by a confidence score Δ*E*/(*n* +1), where Δ*E* is the energy gap (Fig. 1E), and *n* is the number of lower-energy matches discarded in assignments made previously, within that species and at that iteration (Fig. S9). All the assigned HK-RR pairs are then ranked in order of decreasing confidence score.
5. **Incrementation of the CA**. At each iteration *n* > 1, the (*n* – 1) *N*_increment_ assigned HK-RR pairs that had the highest confidence scores at iteration *n* – 1 are included in the CA. In the presence of a gold standard set, the *N*_start_ gold standard HK-RR pairs are also included in the CA. Without a gold standard set, the initial CA is built by randomly pairing each HK of the dataset to an RR from the same species, and for *n* > 1, the CA only contains the above-mentioned (n – 1)*N*_increment_ assigned pairs. Once the new CA is constructed, the iteration is completed, and the next one can start.

## ACKNOWLEDGMENTS

We thank Mohamed Barakat and Philippe Ortet for sharing and discussing specifically-formatted datasets built from the P2CS database. AFB acknowledges support from the Human Frontier Science Program. This research was supported in part by National Institutes of Health Grant R01-GM082938 (AFB and NSW) and by National Science Foundation Grant PHY1305525 (NSW).

## AUTHOR CONTRIBUTIONS

A.F.B., R.S.D., L.J.C. and N.S.W. designed research, A.F.B., L.J.C. and N.S.W. performed research, analyzed data, and wrote the paper.

## NOTE

While submitting this manuscript, we learned that a group led by Andrea Pagnani and Martin Weigt is preparing a related paper on predicting paralog pairs.

## SUPPORTING INFORMATION

EXTENDED MATERIALS AND METHODS

### I DATASET CONSTRUCTION

#### A Complete dataset

Our dataset is based on the online database P2CS (http://www.p2cs.org/) [33, 34], which includes two-component-system proteins from all fully-sequenced prokaryotic genomes. In the construction of P2CS, these proteins were identified by searching genomes for two-component system domains from the Pfam (http://pfam.xfam.org/) and SMART (http://smart.embl-heidelberg.de/) libraries. We kept only chromosome-encoded proteins, due to strong variability in plasmid presence. We also excluded the hybrid and unorthodox proteins, which involve both HK and RR domains in the same protein, since the energetics of partnering is different and often less constraining for such proteins [13]. In HKs, there are different variants of the domain around the N-terminal Histidine-containing phosphoacceptor site, which includes the region that interacts with RRs. These domains are classified into several different Pfam domain families, all members of the His_Kinase_A domain clan (CL0025). In order to reliably align all HK sequences together, we chose to focus on only one of these Pfam domain families, HisKA (PF00512). Proteins containing a HisKA domain account for the majority (64%) of all chromosome-encoded, non-hybrid, orthodox HKs in P2CS.

Proteins in P2CS are annotated based on genetic organization [34]. As our aim was to benchmark our method on known, specific interaction partners, we only considered HKs and RRs in pairs, i.e. encoded by adjacent genes. Note that 67% of all chromosome-encoded, non-hybrid, orthodox HKs in P2CS are from such pairs. Suppressing the (rare) HKs with multiple HisKA domains and RRs with multiple response_reg domains for which the pairing of domains is ambiguous, this yields 23,632 distinct pairs (that differ at least by their sequences or by their species). Discarding those from species with only one such pair, for which HK-RR pairing is trivial, we finally obtained a complete curated dataset of 23,424 HK-RR pairs. Grouping together sequences with mean Hamming distance per site < 0.3 (i.e. with 70% sequence identity or more) to estimate sequence diversity yields an effective number of HK-RR pairs *M*_eff_ = 5391 in the complete dataset.

These 23,424 HK-RR pairs are from 2102 different species, with numbers of pairs per species ranging from 2 to 41, with mean 〈mp〉 = 11.1. The distribution of the number of pairs per species in our complete dataset is shown in Fig. S6A.

#### B Standard dataset

In most of our work, we focused on a smaller “standard dataset” extracted from this complete dataset, both because protein families that possess as many members as the HKs and RRs are atypical, and in view of computational time constraints. Note, however, that our IPA was used to make predictions on the complete dataset, yielding a striking 0.93 final TP fraction (see Fig. 5B).

Our standard dataset was constructed by picking species randomly, and it comprises 5064 pairs from 459 species, with an average number of pairs per species 〈*m_p_*〉 = 11.0, which is very close to that of the complete dataset (see Fig. S6A for the distributions of the number of pairs per species). Grouping together sequences with mean Hamming distance per site < 0.3 to estimate sequence diversity yields an effective number of HK-RR pairs *M*_eff_ = 2091 in the standard dataset.

#### C Multiple sequence alignment

All HKs in our dataset were aligned to the profile hidden Markov model (HMM) representing the Pfam HisKA domain (PF00512) using the hmmalign tool from HMMER (http://hmmer.org/) [35]. Similarly, all RRs were aligned to the profile HMM representing the Pfam Response_reg domain (PF00072). The aligned sequences of each HK were then concatenated to those of their RR partner, yielding a concatenated multiple sequence alignment. The length of each concatenated sequence is *L* = 176 amino acids, among which the *L*_HK_ = 64 first amino acids are from the HK, and the remaining 112 amino acids are from the RR. The full length of these sequences was kept throughout.

### II STATISTICS OF THE CONCATENATED ALIGNMENT (CA)

Let us consider a CA of paired HK-RR sequences. At each site *i* ∊ {1,.., *L*}, where *L* is the number of amino-acid sites, a given concatenated sequence can feature any amino acid (denoted by *α* with *α* ∊ {1,.., 20}), or a gap (denoted by *α* = 21), yielding 21 possible states *α* for each site *i*.

To describe the statistics of the alignment, we only employ the single-site frequencies of occurrence of each state *α* at each site *i*, denoted by 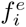(*a*), and the two-site frequencies of occurrence of each ordered pair of states (*α*,*β*) at each ordered pair of sites (*i,j*), denoted by 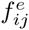(*α*,*β*) [7]. The raw empirical frequencies, obtained by counting the sequences where given residues occur at given sites and dividing by the number *M* of sequences in the CA, are subject to sampling bias, due to phylogeny and to the choice of species that are sequenced [9, 28]. Hence, to define 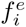 and 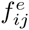, we use a standard correction that re-weights “neighboring” concatenated sequences with mean Hamming distance per site < 0.3. The value of this similarity threshold is arbitrary, but we confirmed that our results depend very weakly on this choice, even when taking the threshold down to zero (data not shown). The weight associated to a given concatenated sequence a is 1/*m*_a_, where *m*_a_ is the number of neighbors of a within the threshold [9, 15, 28]. This allows one to define an effective sequence number *M*_eff_ via

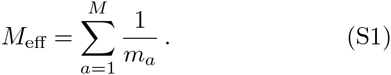

To avoid issues due to the finite size of the CA, such as amino acids that never appear at some sites, which would present mathematical difficulties, e.g. a non-invertible correlation matrix and diverging couplings [28], we introduce pseudocounts via a parameter Λ [7, 9, 15, 28]. The corrected one-site frequencies *f_i_* become

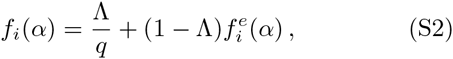

where *q* = 21 is the number of states (i.e. of amino acids, including gaps) per site. Similarly, the corrected two-site frequencies *f_ij_* become

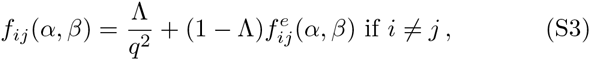

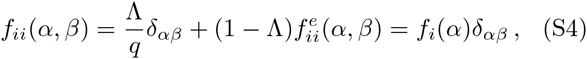

where *δ_αβ_* = 1 if *α* = *β* and 0 otherwise. These pseudo-count corrections are uniform (i.e. they have the same weight 1/*q* on all amino-acid states), and their importance relative to the raw empirical frequencies can be tuned through the parameter Λ. In practice, we take Λ = 0.5, which has been shown to be a satisfactory choice [9, 28]. Note that the correspondence of Λ with the parameter λ in Refs. [9, 15, 28] is obtained by setting Λ = λ/(λ + *M*_eff_).

From these quantities, we define the two-point correlations

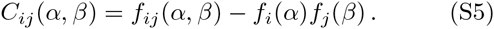

### III MAXIMUM ENTROPY MODEL

#### A Formulation

The maximum entropy principle [8] yields the following form for the least-structured global (*L*-point) probability distribution *P* of sequences consistent with the empirical one- and two-point statistics of the CA:

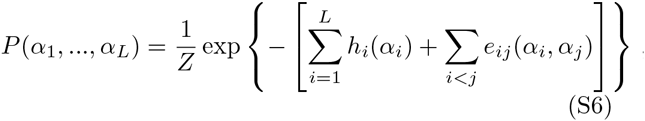

where *Z* is a normalization constant. Each one-body term *h_i_* is known as the field at site *i*, and each two-body interaction term *e_ij_* is known as the (direct) coupling between sites *i* and *j*. The fields *h_i_* and the couplings *e_ij_* are determined by imposing that the probability distribution *P* be consistent with the empirical one- and two-point frequencies *f_i_* and *f_ij_*:

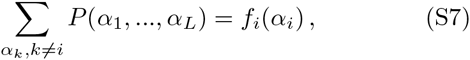

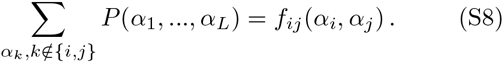

Such pairwise interaction maximum entropy models have proved very successful in various fields (see e.g. Refs. [12, 14, 20-25]), including the prediction of protein structures and inter-protein contacts from multiple sequence alignments (see e.g. Refs. [7, 9, 28]). In particular, high couplings *e_ij_* are better predictors of real contacts in proteins than high correlations *C_ij_*, because the *e_ij_* represent minimal direct couplings between amino acids, while high *C_ij_* can arise from indirect effects [7, 9, 28].

#### B Inference of the parameters

Eqs. S7 and S8 alone do not uniquely define all the fields *h_i_* (*α*) and couplings *e_ij_*(*α,β*) with 1 ≤ *i* < *j* ≤ *L* involved in Eq. S6, which amount to *Lq* + *L*(*L* – 1)*q*^2^/2 parameters, where *q* = 21 is the number of amino-acid states *α*. Indeed, while the number of equations in Eqs. S7 and S8 is the same as that of the empirical frequencies, the latter are not all independent. The two-site frequencies are symmetric (*f_ij_* (*α*,*β*) = *f_ji_* (*β*,*α*)) and consistent with the one-site frequencies (*f_ii_* (*α*, *β*) = *f_i_* (*α*)*δ_αβ_*; Σ*_β_ f_ij_* (*α*,*β*) = *f_i_* (*α*); and Σ*_α_ f_ij_* (*α*,*β*) = *f_j_* (*β*)), which sum to one (Σ*_α_ f_i_* (*α*) = 1). All these constraints reduce the number of independent variables among the one- and two-site frequencies, and thus of independent equations, to *L*(*q* – 1) + *L*(*L* – 1)(*q* – 1)^2^/2 [7, 28]. This yields a gauge degree of freedom in the determination of the fields and couplings from Eqs. S7 and S8. Given the number of independent equations, one possible gauge choice is to set to zero the fields and couplings for one given state, e.g. state *q* (the gap) [9, 28]: *h_i_*(*q*) = 0 and, for all *α*,

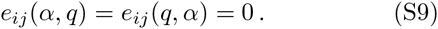

Determining the remaining fields *h_i_* and the couplings *e_ij_* from Eqs. S7 and S8 is difficult, and various approximations have been developed to solve this problem. Following Refs [9, 28], we use the mean-field or small-coupling approximation, which was introduced in Ref. [36] for the Ising spin-glass model. In this approximation, for *i* ≠ *j* and *α*, *β* < *q*, the couplings are given by *e_ij_* (*α*, *β*) = 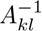, where *A* is a (*q* – 1)*L* × (*q* – 1)*L* correlation matrix: *A_kl_* = *C_ij_* (*α*, *β*), where *k* = (*q* – 1)(*i* – 1) + *α* and *l* = (*q* – 1)(*j* – 1) + *β* [30]. This can be summarized as

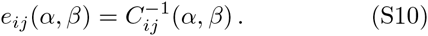

Together, Eqs. S9 and S10 yield all the couplings. Note that the couplings are symmetric (*e_ij_* (*α*,*β*) = *e_ji_*(*β*,*α*)) since the correlations are.

This simple mean-field approximation has been used with success for protein structure prediction [9, 28]. (More sophisticated approximations typically improve performance by less than ten percent [16, 30].) Moreover, this approximation is computationally fast, since it only requires the inversion of a (20*L*) × (20*L*) correlation matrix. Computational rapidity is a considerable asset for our purpose, given that the IPA performs better with smaller increment step size *N*_increment_ (see Fig. 3), i.e. with more iterations, and that the couplings *e_ij_* are computed at each iteration. This approximation also enabled us to use the full-length sequences to infer couplings, without needing to restrict to a subset of amino-acid sites as in some other works using more sophisticated approximations [7, 15]. Interestingly, we found that using full-length sequences contributed to increasing the TP fraction in our framework (data not shown).

#### C Gauge choice

Qualitatively, the gauge degree of freedom means that contributions to the effective energy of the system (which is minus the argument of the exponential in Eq. S6) can be shifted between the fields and the couplings [7]. Since our focus is on interactions, we do not want the couplings to include contributions that can be accounted for by the (one-body) fields [37]. The zero-sum (or Ising) gauge, where the couplings satisfy

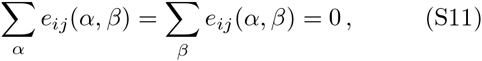

minimizes the Frobenius norms of the couplings

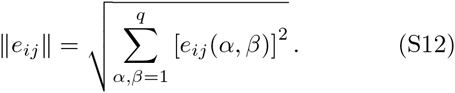

Hence, the zero-sum gauge attributes the smallest possible fraction of the energy in Eq. S6 to the couplings, and the largest possible fraction to the fields [7, 30]. Furthermore, when employing this gauge, the Frobenius norm has proved to be a successful predictor of contacts in proteins [16, 30]. In particular, within the mean-field approximation Eq. S10, the use of the Frobenius norm (with an average-product correction) improves over the results obtained using direct information [30].

Thus, after calculating the couplings as described above, we change the gauge from the one defined in Eq. S9 to the one defined in Eq. S11, by replacing each coupling *e_ij_*(*α,β*) by

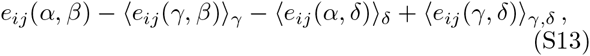

where 〈.〉_γ_ denotes an average over γ ϵ {1,..., *q*} [30].

Note that in Fig. 4, we use the Frobenius norm without the average-product correction [30]. With this correction, implemented by averaging within single proteins [17], we obtained similar results (data not shown). However, with the correction, final performance appeared slightly lower, but training was visible slightly earlier in the IPA, likely because phylogenetic bias and insufficient sampling have a stronger impact at early stages.

### IV ITERATIVE PAIRING ALGORITHM (IPA)

The main steps of the IPA are shown in Fig. 1C. Here, we describe each of these steps in detail, after explaining how the CA is constructed for the very first iteration.

#### Initialization of the CA

##### Starting from a gold standard set of HK-RR pairs

Since the gold standard pairs are considered as known specific interaction partners, i.e. “reference” pairs, the CA for the first iteration of the IPA corresponds to this gold standard set. In subsequent iterations, the pairs from the gold standard set are *always kept* in the CA, and additional pairs from the rest of the dataset, those with the highest confidence scores (see below), are included in the CA.

##### Starting from random pairings

In the absence of a gold standard set, each HK of the dataset is randomly paired with an RR from its species, using a random permutation in each species. All these *M* pairs (where *M* represents the total number of HKs, or, equivalently, RRs, in the dataset) are included in the CA for the first iteration of the IPA. Hence, this initial CA contains a mixture of correct and incorrect pairs, with one correct pair per species on average. At the second iteration, the CA is built only from the *N*_increment_ HK-RR pairs with the highest confidence scores obtained from this first iteration.

There are other ways to initialize the CA in the absence of a gold standard set. We varied the number of pairs included at the second iteration (which is *N*_increment_ in the above scheme), and we also tried including in the first CA all possible HK-RR pairs from the species with few pairs (for which this exhaustive pairing yields a larger proportion of true pairs). These variants did not significantly increase the final TP fraction (data not shown). Moreover, the random initialization of the CA can be exploited to increase the TP fraction (Figs. 6 and S7), which would be impossible for exhaustive initializations.

Now that we have described the initial construction of the CA, we describe each step of an iteration of the IPA (Fig. 1C).

#### Step 1: Correlations

At each iteration, the empirical one- and two-body frequencies are computed for the CA, using the re-weighting of neighbor sequences and the pseudocount correction described above (see Eqs. S1-S4). The empirical correlations *C_ij_* are then deduced using Eq. S5.

#### Step 2: Direct couplings

The direct couplings in the pairwise maximum entropy model of the CA are inferred from the empirical correlations of the CA using Eqs. S9 and S10. The gauge is then changed to the zero-sum gauge (Eq. S11) by making the transformation in Eq. S13.

#### Step 3: Interaction energies for all possible HK-RR pairs

The interaction energy *E* of each possible HK-RR pair within each species of the dataset is calculated by summing the appropriate direct couplings:

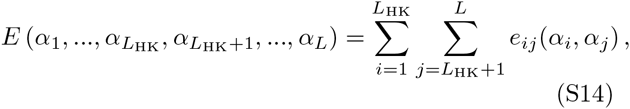

where *L*_HK_ denotes the length (i.e. the number of amino-acid sites) of the HK sequence and *L* that of concatenated HK-RR sequence. Note that this HK-RR interaction energy only involves the inter-molecular couplings (*i* ≤ *L*_HK_ and *j* > *L*_HK_; the case *i* > *L*_HK_ and *j* ≤ *L*_HK_ does not need to be considered as the couplings are symmetric).

#### Step 4: HK-RR pair assignments and ranking by energy gap

##### HK-RR pair assignments

In each separate species, the pair with the lowest interaction energy is selected first, and the HK and RR from this pair are removed from further consideration, since we assume one-to-one HK-RR matches (see Fig. 1D). Then, the next pair with the lowest energy is chosen, and the process is repeated until all HKs and RRs are paired.

#### Scoring by gap

Each assigned HK-RR pair is scored at the time of assignment by Δ*E*/(*n*+1), where Δ*E* is the energy gap (see Fig. 1E), and *n* is the number of lower-energy matches discarded in assignments made previously (within that species and at that iteration). Qualitatively, the larger the discrepancy between the best match and the next best one (i.e. the larger the energy gap), and the fewer the rejected better candidates (i.e. the smaller *n*), the more reliable we expect the assignment to be.

More precisely, Δ*E*_RR_ = *E*_RR,2_ — *E*_RR,1_ > 0 is computed for the RR involved as minus the difference of the interaction energy *E*_RR,1_ of this RR with its assigned partner (i.e. the “best” HK, which yields the lowest interaction energy with this RR, *among the HKs that are still unpaired*) and that *E*_RR,2_ with the second-best HK *among the HKs that are still unpaired*. Meanwhile, *n*_RR_ is the number of HKs of that species that had lower interaction energies with this RR than the assigned partner, but that have been eliminated previously in that iteration’s pairing process, because they were paired to other RRs with a lower interaction energy. A schematic example is shown on Fig. S9A. Similarly, the value of Δ*E*_HK_ and of *n*_HK_ are calculated for the HK involved in the assigned pair. Finally, the lowest score among the two obtained is kept:

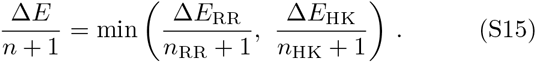

We have chosen to divide the energy gap Δ*E* by *n* + 1 in order to penalize the HK-RR pairs made after better candidates were discarded, even if their current gap among remaining candidates appears large, as illustrated by the second assignment in Fig. S9A. However, one could consider other definitions of confidence scores, such as Δ*E*/(*n* + 1)^α^, where *α* is a parameter. In Fig. S9B, we show that our confidence score significantly improves TP fraction over the raw energy gap Δ*E*, and that *α* = 1 yields an optimal TP fraction.

This definition of the confidence score leaves an ambiguity for the last assigned pair of each species, since there is no remaining second-best match to define the energy gap. We have chosen to assign to this pair a confidence score equal to the lowest other one within the species, given that this pair, made by default, should not be deemed more reliable than any other pair in the species. Another ambiguity exists in the (rare) case where several pairs have the exact same interaction energy. In this case, we chose to make one assignment between the equivalent matches, and to leave the other equivalent HKs and/or RRs to be paired later. We checked that the impact of this choice on final results is very small.

#### Ranking of pairs

Once all the HK-RR pairs are assigned and scored, we rank them in order of decreasing confidence score.

#### Step 5: Incrementation of the CA

The ranking of the HK-RR pairs based on the confidence score is used to pick the pairs that are included in the CA at the next iteration. Pairs with a high confidence score are more likely to be correct because there was less ambiguity in the assignment. The number of pairs in the CA is increased by *N*_increment_ at each iteration, and the IPA is run until all the HKs and RRs in the dataset are in the CA (in the last iteration, all pairs assigned at the second to last iteration are included in the CA).

#### Starting from a gold standard set of HK-RR pairs

The *N*_start_ gold standard pairs are kept in the CA in all iterations and the HKs and RRs involved in these pairs are not paired or scored by the IPA. The HKs and RRs from all the other pairs in the CA are always repaired and re-scored at each iteration, and all the non-gold-standard pairs of the CA are re-picked based on their confidence score. In other words, at the first iteration, the CA only contains the *N*_start_ gold standard pairs. Then, for any iteration number *n* > 1, it contains these exact same *N*_start_ gold standard pairs, plus the (*n* – 1)*N*_increment_ assigned HK-RR pairs that had the highest confidence scores at iteration number *n* – 1.

#### Starting from random pairings

In the absence of a gold standard set, all M HKs and RRs in the dataset are paired and scored at each iteration, and all the pairs of the CA are fully re-picked at each iteration based on the confidence score. The first iteration is special, since the CA is made of *M* random within-species HK-RR pairs (see above, “Initialization of the CA”). Then, for any iteration number *n* > 1, the CA contains the (*n* – 1)*N*_increment_ assigned HK-RR pairs that had the highest confidence scores at iteration number *n* – 1.

Once the new CA is constructed, the iteration is completed, and the next one can start with Step 1, the computation of the empirical correlations in this CA.

**FIG. S1.**
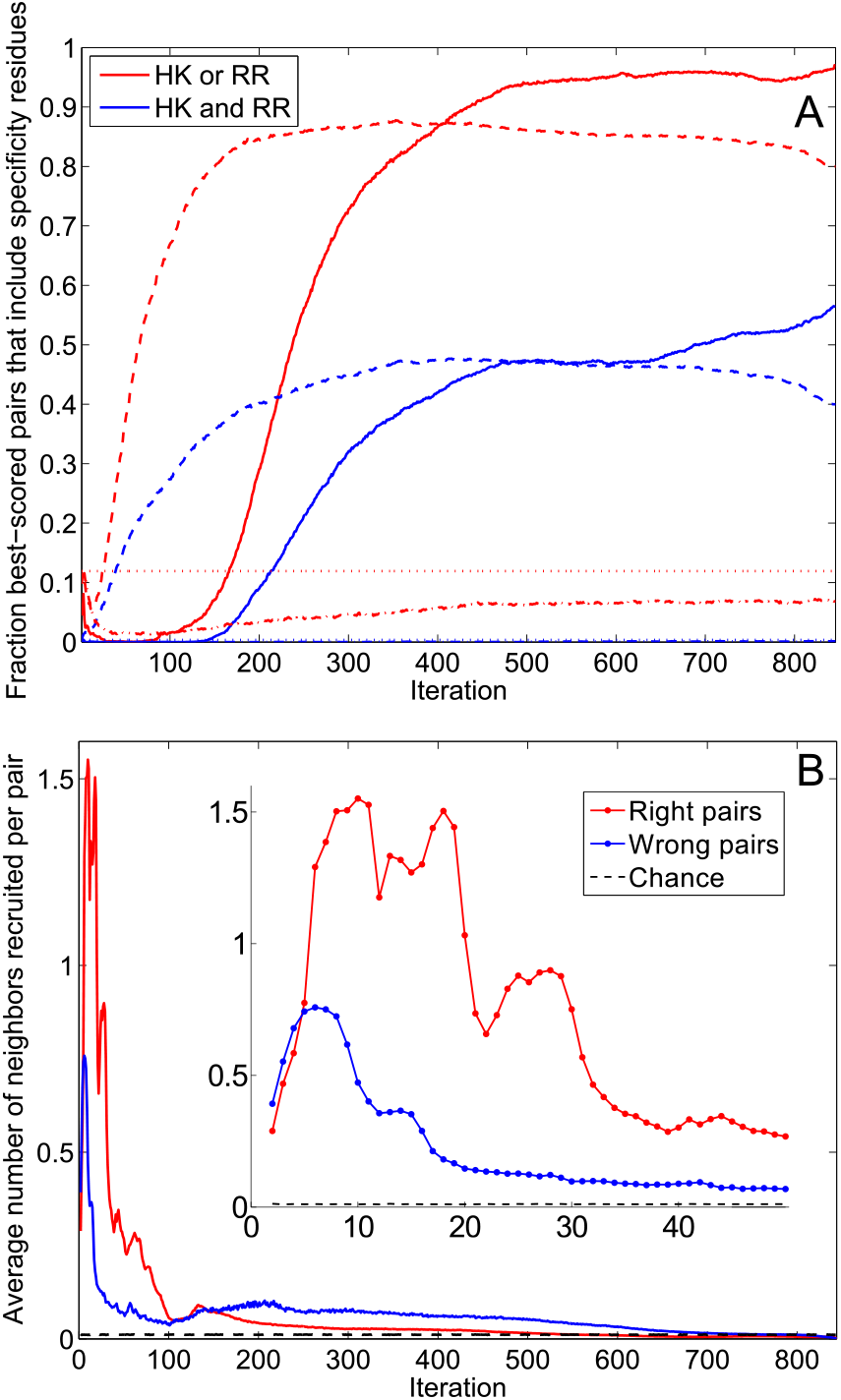
Evolution of the coupling matrix and of the concatenated alignment (CA) during the IPA. (A) Training of the coupling matrix. As in Fig. 4A, pairs comprised of an HK residue site and an RR residue site are scored by the Frobenius norm (i.e. the square root of the summed squares) of the couplings involving all possible residue types at these two sites. The 10 best-scored pairs are compared to the main specificity residues determined experimentally in Refs. [4, 38–40] (5 HK residues, T267, A268, A271, Y272, and T275 in the sequence of *T*. *maritima* HK853, and 5 RR residues, V13, L14, I17, N20, and F21 in the sequence of *T*. *maritima* RR468 [39]). Solid curves: Fraction of the 10 best-scored residue pairs that include HK and/or RR specificity residues versus the iteration number in the IPA. Dashed curves: Ideal case, where at each iteration *N*_increment_ randomly-selected correct HK-RR pairs are added to the CA. Dash-dotted curves: Case where random HK-RR pairs are added to the CA. Dotted lines: Overall fraction of residue pairs that include specificity residues. (B) Neighbor recruitment. Average number of neighbors an HK-RR pair of the CA has among the new HK-RR pairs of the next CA versus iteration number. Two pairs are considered neighbors if the mean Hamming distance per site between the two HKs and between the two RRs are both < 0.3. Dashed line: Null model – at each iteration, *N*_increment_ new correct HK-RR pairs are chosen at random and added to the CA. Inset: Expanded view of the first 50 iterations. In both panels, the IPA is performed on the standard dataset with *N*_increment_ = 6. In panel A (resp. B), data is averaged over 500 (resp. 5193) replicates that differ in their initial random pairings.

**FIG. S2.**
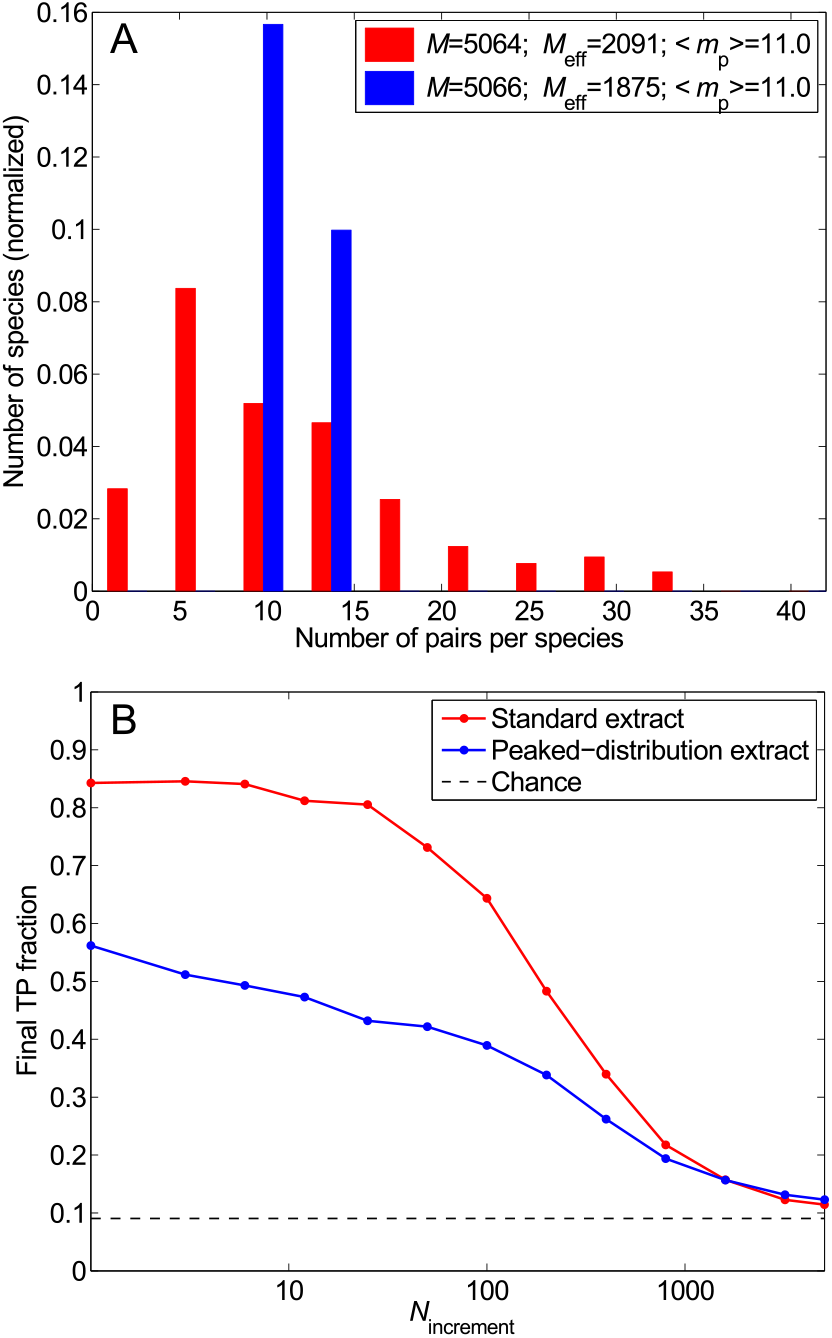
Impact of the distribution of the number of HK-RR pairs per species. (A) Distribution of the number of pairs per species in two different datasets: the standard one (red) and one with the same total number of HK-RR pairs *M* and the same mean number of pairs per species 〈*m_p_*〉, but with a more strongly peaked distribution (blue). (B) Final TP fraction versus *N*_increment_ for the two datasets described in (A). All results are averaged over 50 replicates that differ in their initial random pairings. Dashed line: Average TP fraction obtained for random HK-RR pairings.

**FIG. S3.**
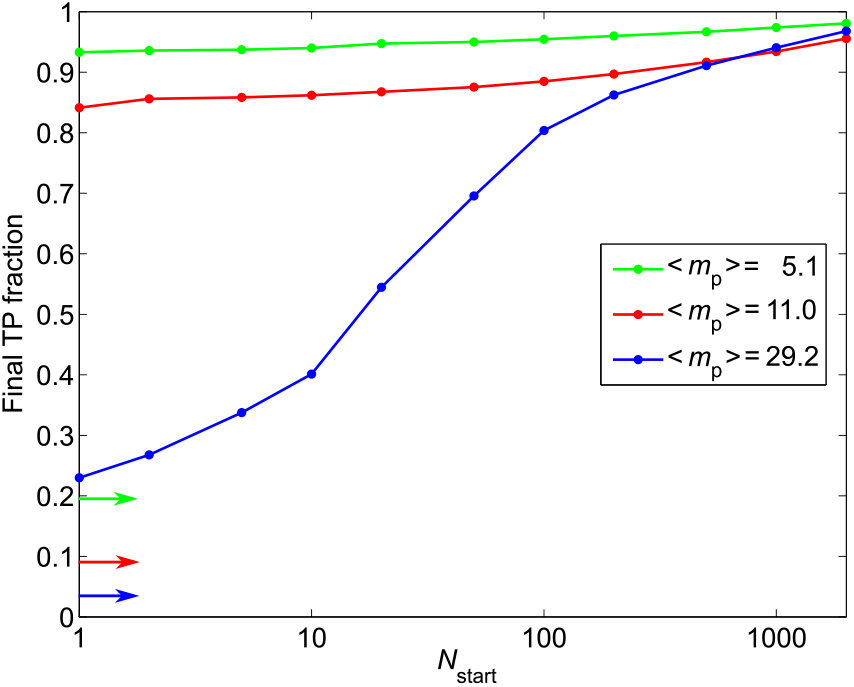
Impact of the number of HK-RR pairs per species, and of the total number of HK-RR pairs in the dataset: starting from a gold standard set. Final TP fraction versus *N*_start_ for the three datasets with different distributions of the number of pairs per species yielding different means 〈*m_p_*〉 presented in Fig. 5A. Colored arrows indicate the average TP fractions obtained for random HK-RR pairings in each dataset. The IPA is performed on the standard dataset with *N*_increment_ = 6. All results are averaged over 50 replicates that differ by the random choice of pairs in the gold standard set.

**FIG. S4.**
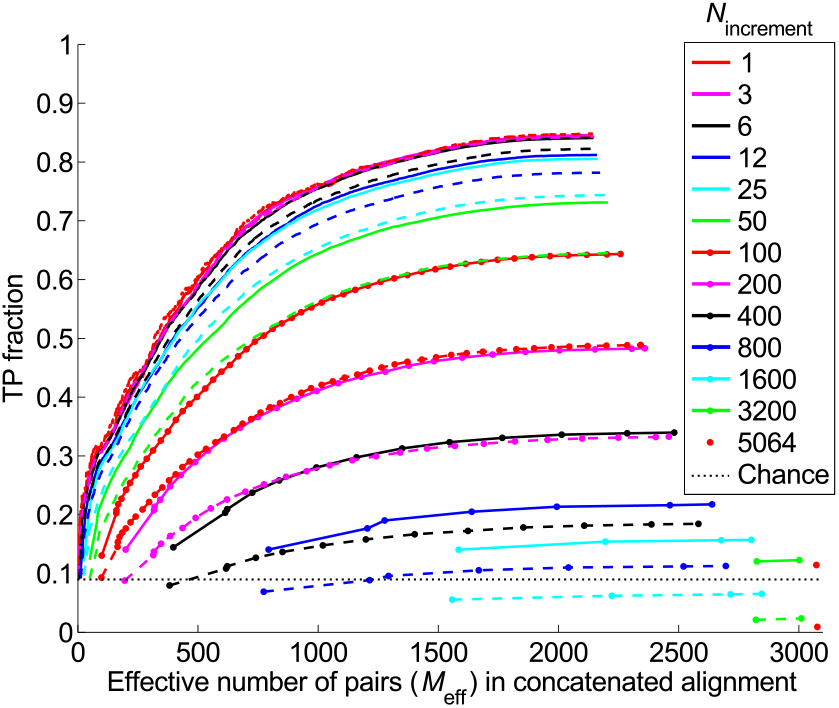
Impact of the initial correct pairs. TP fraction versus effective number of HK-RR pairs (*M*_eff_) in the concatenated alignment during iterations of the IPA, for different values of *N*_increment_. Solid curves: Starting from random pairings (data also shown in Fig. 3). Dashed curves: Starting from random pairings with no initial correct pair (the color and symbol codes are the same as for the solid curves). The standard dataset is used. All results are averaged over 50 replicates that differ in their initial random pairings. Dotted line: Average TP fraction obtained for random HK-RR pairings.

**FIG. S5.**
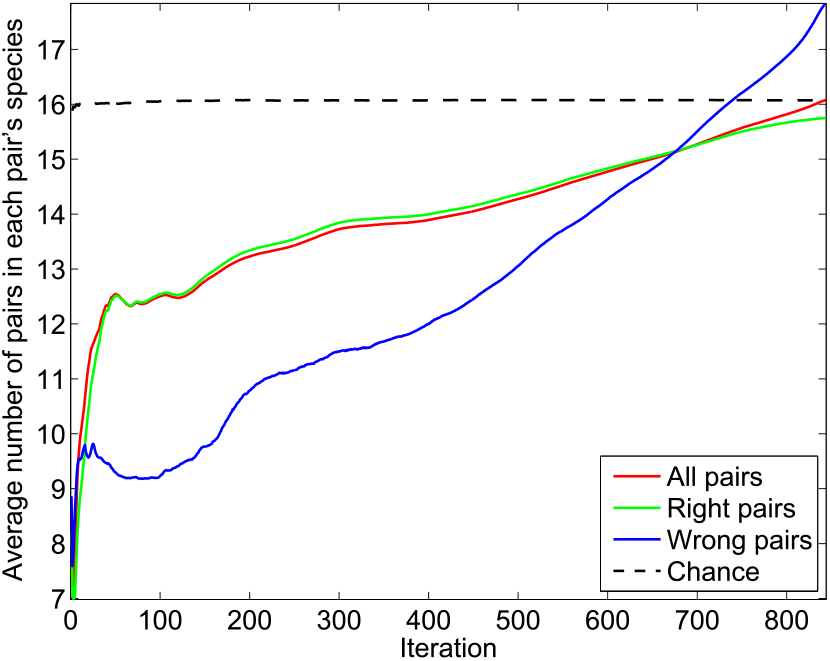
Evolution of the concatenated alignment (CA) during the IPA. Average number of HK-RR pairs present in the species to which the pairs of the CA belong versus iteration number. The IPA is performed on the standard dataset, with *N*_increment_ = 6, and all data is averaged over 5193 replicates that differ in their initial random pairings. Dashed line: At each iteration, 6 new correct HK-RR pairs are chosen at random and added to the CA. This chance result just matches the average number of pairs in a pair’s species: 16.1. Note that this number is different from the above-discussed average number of pairs per species 〈*m_p_*〉, which is 11.0 in the standard dataset (because the average over the pairs is not the same as the average over the species).

**FIG. S6.**
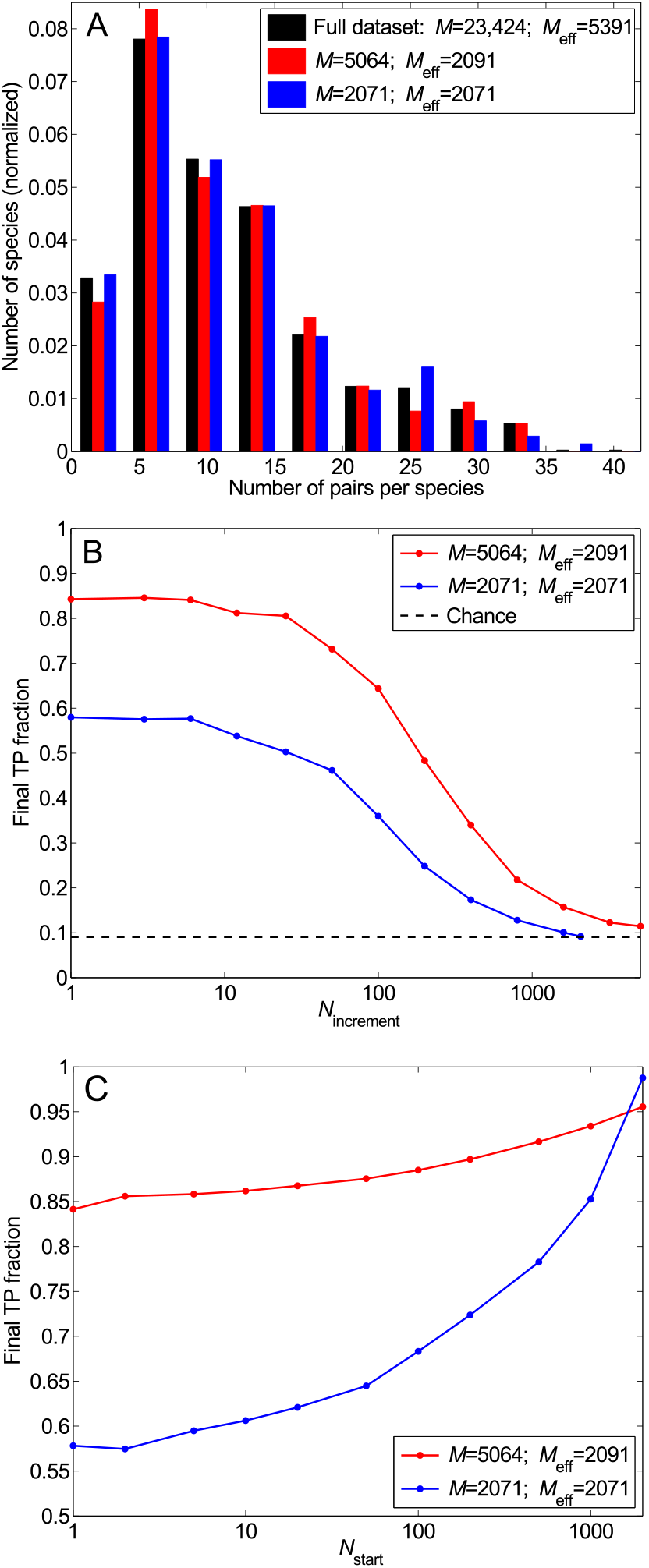
Impact of sequence similarity in the dataset. (A) Distribution of the number of pairs per species in the complete dataset (black) and in two smaller selected datasets each with the same effective number of HK-RR pairs *M*_eff_: the standard one (red) and one where similar sequences have been suppressed such that no two pairs have a mean Hamming distance per site < 0.3 (blue). (B) Final TP fraction versus *N*_increment_ for the two selected datasets described in (A), starting from random pairings. Dashed line: Average TP fraction obtained for random HK-RR pairings. (C) Starting from a gold standard set. Final TP fraction versus *N*_start_ for the two selected datasets presented in (A), with *N*_increment_ = 6. In (B) and (C), all results are averaged over 50 replicates.

**FIG. S7.**
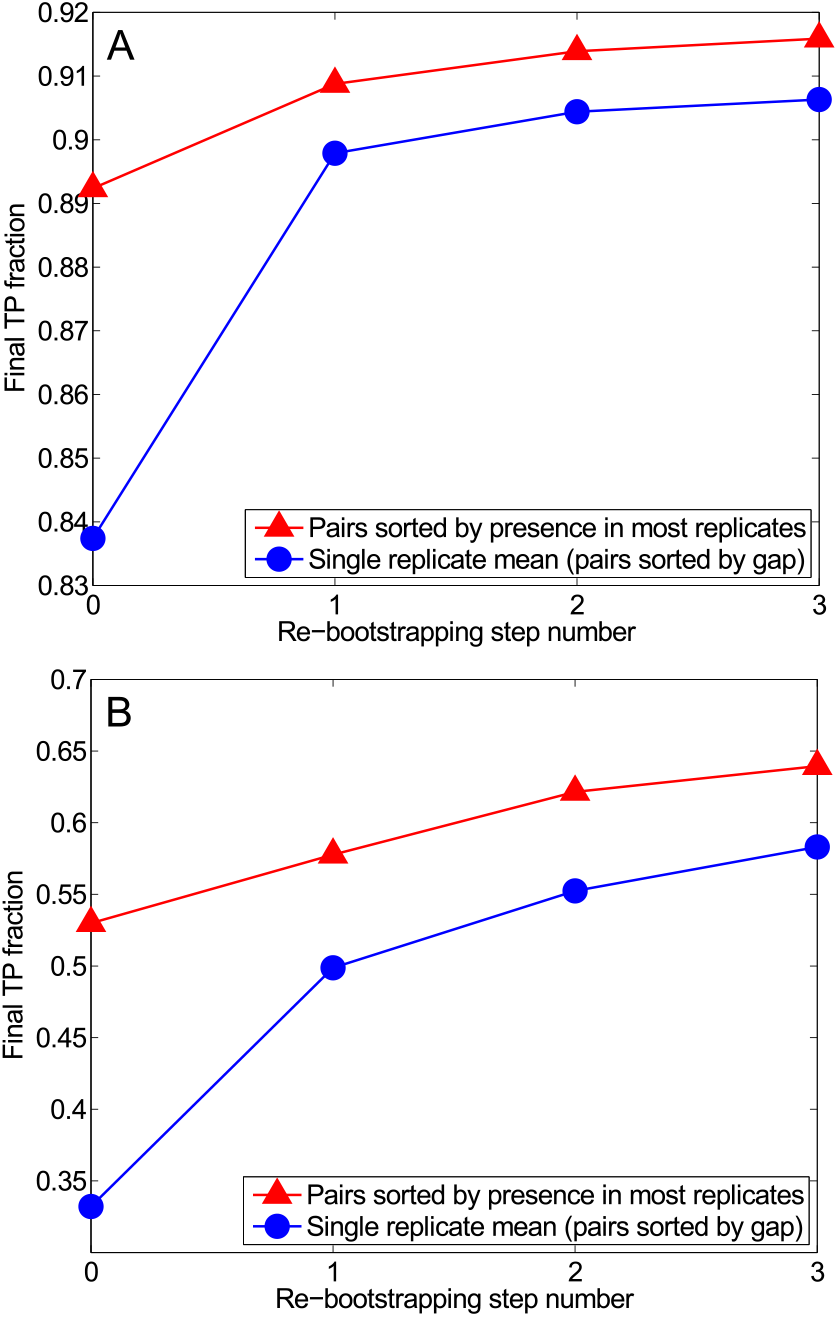
Re-bootstrapping: exploiting the high TP fraction of the HK-RR pairs predicted to be correct in most replicates of the IPA, which differ in their initial random pairings. (A) Re-bootstrapping on the standard dataset (*M* = 5064 HK-RR pairs). The final TP fraction is plotted versus re-bootstrapping step number. Step 0 corresponds to the standard bootstrapping procedure described in the main text (IPA starting from random pairings, see Fig. 6). 500 replicates are computed. We then take as a gold standard set 1000 HK-RR pairs chosen randomly among those predicted to be correct in more than 50% of replicates. These pairs are chosen with probability equal to the fraction of replicates in which they are predicted to be true. The IPA is then performed again starting from such gold standard sets. The process is then iterated. Here, 50 replicates were computed for steps 1, 2, and 3. The average final TP fraction is plotted (blue curve), as well as the TP fraction for the best *M* = 5064 pairs ranked by the fraction of replicates in which they are predicted to be true (red curve, see Fig. 6). Here, *N*_increment_ = 6. (B) Re-bootstrapping on a smaller dataset with *M* = 502 HK-RR pairs from 40 species (mean number of pairs per species 〈*m_p_*〉 = 12.6). The process is the same as in (A), but here, at each re-bootstrapping step, we take as a gold standard set 200 HK-RR pairs chosen randomly among those predicted to be true in more than 25% of replicates at the previous step, and *N*_increment_ = 1.

**FIG. S8.**
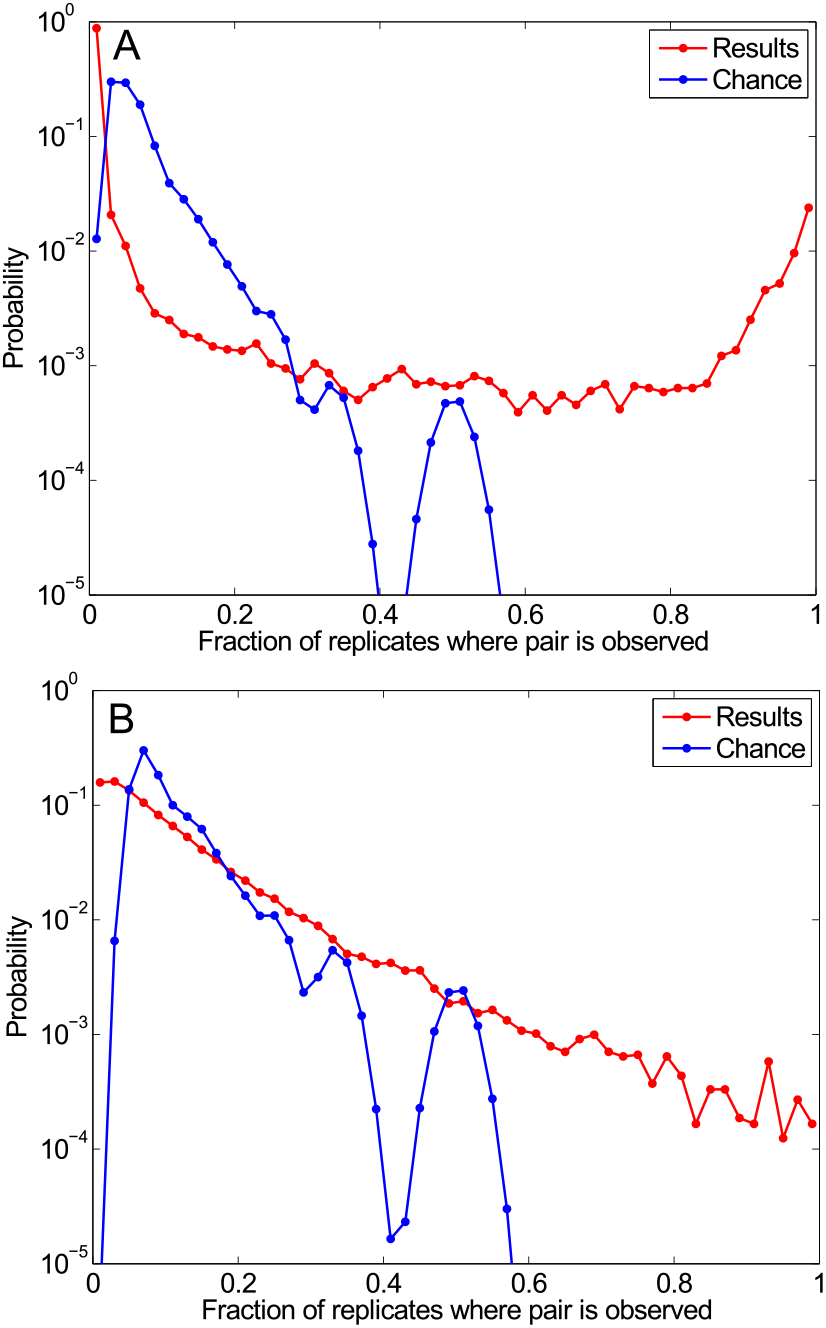
Distribution of the fraction of replicates *f_r_* in which each pair is predicted to be true. (A) Standard dataset. Red curve: For each possible HK-RR pair (within each species), *f_r_* is estimated over 500 replicates of the IPA that differ in their initial random pairings, with *N*_increment_ = 6 (same data as in Fig. 6). Blue curve: Random HK-RR pairings within each species of the standard dataset. (B) Dataset with no correct pairs. A dataset of the same size as the standard one (*M* = 5062 in practice) that does not include any true HK-RR pairs was constructed. Red curve: Distribution of *f_r_* obtained by applying the IPA to this dataset, exactly as in (A). Blue curve: Random HK-RR pairings within each species of the dataset with no correct pairs. (Note that the blue curve is slightly different from that in (A) due to slight differences in the distribution of the number of pairs per species between the two datasets.) In (A) and (B), the data is binned into 50 equally-spaced bins between *f_r_* = 0 and *f_r_* = 1.

**FIG. S9.**
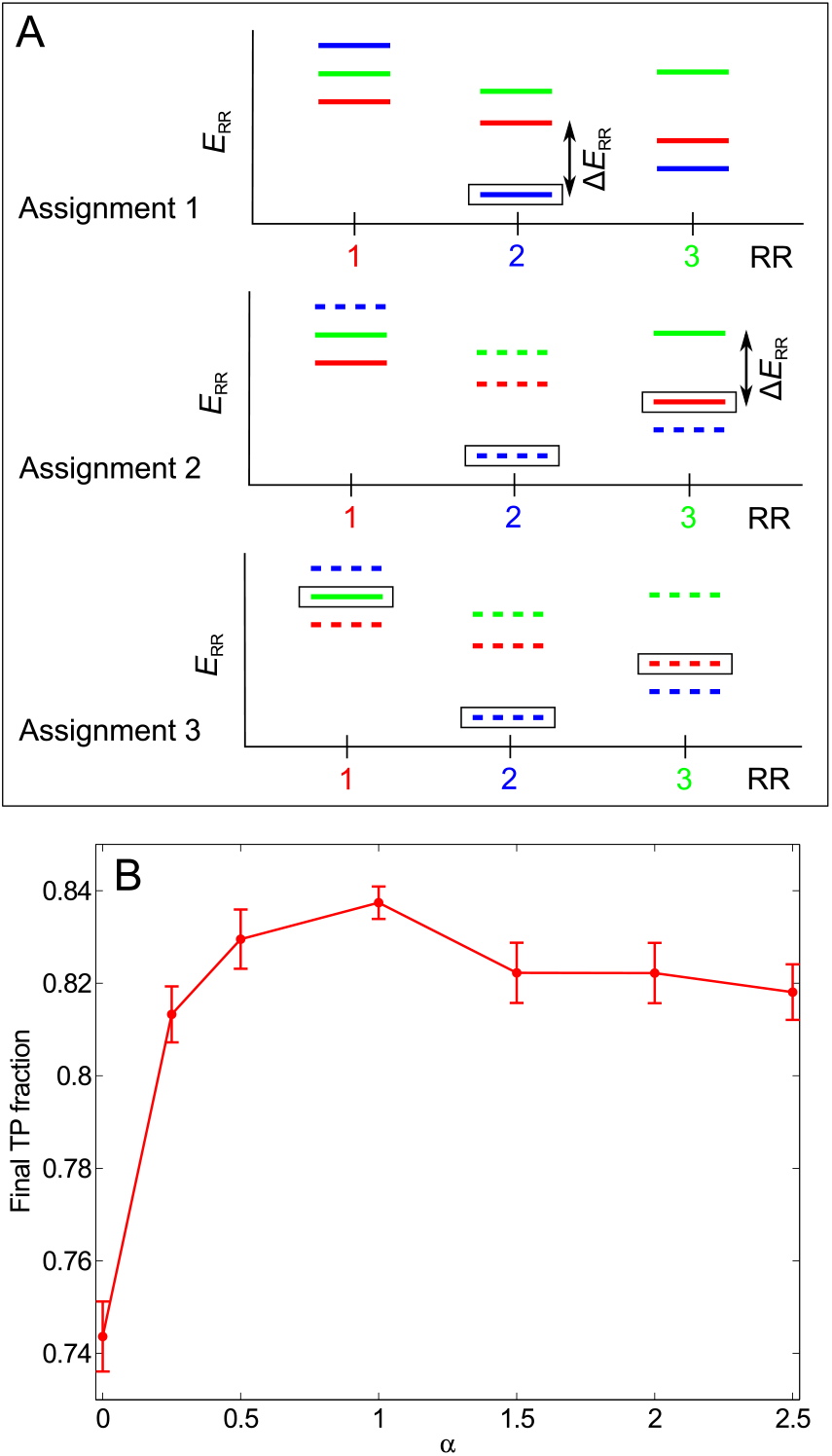
Scoring by gap. (A) Determination of the confidence score of each assigned HK-RR pair in a given iteration of the IPA. In this schematic, we consider a species with three HKs and RRs. In the energy spectra showing the interaction energies for each RR with all three HKs, each color represents a given HK (red: HK 1, partner of RR 1; blue: 2; green: 3). Assignment 1: The pair with the lowest interaction energy (HK 2 - RR 2, boxed) is selected. The energy gap Δ*E*_RR_ is shown. Here *n*_RR_ = 0 since no HK has been removed from consideration yet. Assignment 2: The HK and RR previously paired are removed from further consideration (dashed energy levels). The next pair with the lowest energy (HK 1 - RR 3, boxed) is chosen among the remaining ones. Here *n*_RR_ = 1 since HK 2, which was paired previously, had a lower interaction energy with RR 3 than HK 1. Using the *ad hoc* confidence score Δ*E*_RR_/(*n*_RR_ + 1), this (incorrect) pair is penalized with respect to the (correct) one made in the first assignment, even though their energy gaps are similar. Assignment 3: Only one possible pair remains. It is made, and its confidence score is taken to be equal to the lowest previously calculated confidence score for that species (the second one here). At each HK-RR pair assignment, symmetric confidence scores Δ*E*_HK_/(*n*_HK_ +1) are also calculated from the energy spectra showing the interaction energies for each HK with all three RRs. The final confidence score of a pair, denoted by Δ*E/*(*n*+1), is the smallest of these two scores, i.e. min{Δ*E*_RR_/(*n*_RR_ + 1), Δ*E*_HK_/(*n*_HK_ + 1)}. (B) More generally, in every iteration of the IPA, each predicted HK-RR pair can be scored by Δ*E/*(*n + 1*)^α^, where *α* is a parameter. Red curve: Average final TP fraction obtained versus *α*; error bars: 95% confidence intervals around the mean. The IPA was performed on the standard dataset, with *N*_increment_ = 6. Results are averaged over 200 replicates that differ in their initial random pairings for all *α* except *α* = 1, for which 500 replicates were computed. As we found the highest TP fraction for *α* = 1, all the results elsewhere in the paper were obtained using *α* = 1.

